# A validated neuronal SH-SY5Y platform reveals critical experimental variables for reproducible Aβ1–42 self-assembly neurotoxicity assessment

**DOI:** 10.64898/2026.07.12.738010

**Authors:** Anne-Cécile Van Baelen, Clément Poteaux, Philippe Robin, Xavier Iturrioz, Steven Panek, Norbert Sewald, Denis Servent, Nicolo Tonali

**Affiliations:** Université Paris-Saclay, CEA, Département Médicaments et Technologies pour la Santé (DMTS), SIMoS, 91191 Gif-sur-Yvette (France); Organic and Bioorganic Chemistry (OCIII), Department of Chemistry, Bielefeld University, Universitätsstraße 25, 33615 Bielefeld (Germany)

**Author notes:** Correspondance: Anne-Cécile Van Baelen, Université Paris-Saclay, CEA, Département Médicaments et Technologies pour la Santé (DMTS), SIMoS, 91191 Gif-sur-Yvette (France),; Nicolo Université Paris-Saclay, CEA, Département, Médicaments et Technologies pour la Santé (DMTS), SIMoS, 91191 Gif-sur-Yvette (France).

**Keywords:** SH-SY5Y cells, neuronal differentiation, amyloid β1-42, cytotoxicity

## Abstract

Reliable in vitro evaluation of amyloid-β (Aβ) toxicity is essential for the development of anti-amyloid therapeutics, yet experimental workflows often lack standardization. In our previous work, we established a reproducible protocol for the synthesis, characterization and controlled aggregation of highly pure Aβ1-42. Here, we address the biological component of this variability by evaluating the impact of neuronal differentiation and toxicity assays on Aβ-induced neurotoxicity. SH-SY5Y cells were differentiated using retinoic acid and brain-derived neurotrophic factor, generating a neuron-like phenotype validated by immunofluorescence, gene expression profiling and resistance to H_2_O_2_-induced oxidative stress. Using this characterized model, we investigated the effects of non-aggregated and pre-aggregated Aβ1-42 species on cell viability and transcriptional responses. Strikingly, Aβ toxicity was highly dependent on the aggregation state of the peptide, the differentiation status of the target cells and the viability assay employed. Our results suggest that the lack of standardization in peptide quality, aggregation procedures, neuronal maturation and toxicity assessment represents a major source of variability in the amyloid field. Together, these findings provide a methodological framework to improve the reproducibility and translational relevance of in vitro screening strategies for anti-amyloid therapeutics.

## 1. INTRODUCTION

Alzheimer’s disease (AD) is a major global health issue, affecting almost 60 million people worldwide, and remains an incurable progressive neurodegenerative disorder ^1^. AD is defined by intracellular neurofibrillary tangles of hyperphosphorylated tau and extracellular beta-amyloid (Aβ) peptides plaques, ultimately leading to neuronal loss and brain atrophy ^2^. Among Aβ peptides, Aβ1-42 is the most aggregation-prone, forming soluble oligomers, protofibrils and fibrils ^3^ ^4^. Increasing evidence indicate that soluble species are most closely associated with early neuronal dysfunction, but their precise structural nature remains debated ^5^ ^6^ ^7^ ^8^ ^9^. However, understanding their cytotoxic mechanisms remains challenging mainly due to difficulties in reproducibly controlling Aβ1-42 aggregation, and in limitations of conventional neuronal cell models.

Indeed, the self-assembly of Aβ1-42 is a complex mechanism in which unfolded monomers undergo a transition through transient oligomeric intermediates leading to fibril formation by on-pathway or off-pathway routes ^10^ ^11^ ^12^. Although fibrils were traditionally considered the main pathological hallmark, increasing attention is given to oligomers, which reduce functional monomer availability, and are thought to contribute to synaptic toxicity and neurodegeneration ^4^ ^5^. Mastering this complex process is essential, and it is important to ensure the use of a well-established *in vitro* protocol for controlled aggregation ^13^ ^14^.

The development of novel therapeutic agents targeting amyloid-β aggregation relies heavily on in vitro screening platforms to evaluate the toxicity of Aβ assemblies ^15^. ( However, the vast majority of these studies are still performed using undifferentiated neuroblastoma cell lines, such as SH-SY5Y, which only partially recapitulate the phenotype and physiology of mature human neurons ^16^. Consequently, the biological responses observed in these simplified models may not accurately reflect neuronal susceptibility to Aβ aggregates, raising concerns about the reliability and reproducibility of current in vitro screening strategies for anti-amyloid compounds.

Reliable neuronal cell culture models remain a central challenge. While primary rodent neurons provide high physiological relevance, with authentic morphology and synaptic activity, they are limited by non-human origin and ethical constraints ^17^. Also, human pluripotent stem cells (ESCs and iPSCs) offer a comprehensive model of neurogenesis and are expandable, but require complex, costly protocols that limit widespread and high-throughput use ^18^ ^19^ ^20^. Consequently, there is sustained interest in alternative *in cellulo* models that are cost-effective, reproducible and scalable. The immortalized human neuroblastoma cell lines SH-SY5Y is a widely used neuronal cellular model ^21^ ^22^. SH-SY5Y cells are relatively easy to culture, exhibit unlimited proliferative capacity and grow robustly under standard conditions, but in their undifferentiated state they display only limited neuronal characteristics ^23^. However, these cells retain the capacity for differentiation to induce a more neuronal phenotype, using serum restriction combined with retinoic acid (RA), frequently supplemented with neurotrophic factors such as brain-derived neurotrophic factor (BDNF) or nerve growth factor, promoting neurite outgrowth and the expression of markers associated with mature neurones ^24^ ^25^ ^26^ ^27^ ^28^. Differentiated SH-SY5Y cells thus provide a valuable intermediate model for screening studies, offering enhanced neuronal features, while maintaining lower technical complexity, cost-effectiveness and scalability compared to primary neurons or iPSC-derived systems, but with slightly reduced physiological relevance ^21^.

However, despite this interest, there is still a lack of studies investigating the effects of well-controlled Aβ1-42 aggregates in differentiated SH-SY5Y cells. Most Aβ1-42 toxicity studies are performed on non-differentiated SH-SY5Y cells, especially in the context of amyloid aggregation inhibitors, and their use remains controversial ^29^ ^30^ ^31^. A few studies have nevertheless highlighted a distinct behaviour of Aβ1-42 in differentiated *vs* non-differentiated cells but have not explored the full relationship between toxicity and aggregation state ^32^ ^33^ ^34^. In addition, the differentiation state of SH-SY5Y cells is rarely systematically validated through morphological analysis, gene expression profiling, and functional assays, leaving aspects of their functional maturation incompletely understood. Such validation is essential for establishing robust and relevant *in cellulo* systems, in particular in comparison with iPSCs, to better understand how distinct Aβ1-42 aggregate species influence neurodegenerative mechanisms.

In this context, we propose three independent levels of analysis to validate differentiation methodology that offers a reliable framework for investigating neuronal outcomes in the presence of well controlled Aβ1-42 species. We evaluated the effects of these Aβ1-42 aggregates on (i) cytotoxicity, assessing both membrane integrity and metabolic activity, and (ii) cellular differentiation, including the expression of neuronal markers. This approach enhances the scientific relevance of differentiated SH-SY5Y cultures and broadens their use as a reproducible, accessible, and high-throughput-compatible model for studying neuronal differentiation and the cellular effects of Aβ1-42 aggregates. Rather than simply comparing differentiated and undifferentiated neuronal cells, this work provides a systematic evaluation of the principal experimental variables that influence the interpretation of Aβ toxicity assays. We demonstrated that controlled aggregation protocol, together with a triple-level validated neuronal model is an essential prerequisite for generating reliable and biologically meaningful data during the preclinical evaluation of anti-amyloid therapeutic candidates.

## 2. MATERIALS & METHODS

### 2.1 Cell culture and differentiation

SH-SY5Y cells (CRL-2266 purchased from ATCC) were cultured in a maintenance medium composed of DMEM/F12 (GIBCO 61965059) supplemented with 10% fetal bovine serum (FBS), penicillin (50 U/mL), streptomycin (50 µg/mL) (P/S) and glutaMAX-CTS (0.5 mM), and incubated at 37 °C in a humidified atmosphere containing 5% CO_2_. Culture medium was changed every 2-3 days, and cells were subcultured upon reaching approximately 70-80% confluence using trypsin-EDTA.

For differentiation experiments, cells were seeded at an appropriate density (2×10⁴ cells/cm²) to avoid over confluence during the treatment period. After cell attachment (typically 24 h post-seeding), differentiation was initiated by replacing the maintenance medium with low-serum (1% FBS) differentiation medium containing retinoic acid (RA, 10 µM, Thermo Scientific 302-79-4) and/or brain-derived neurotrophic factor (BDNF, 50 ng/mL, Sigma Aldrich B3795). Differentiation medium was changed every 48 h. Control conditions included treatment with RA alone, BDNF alone, or vehicle.

### 2.2 Immunofluorescence

For immunofluorescence experiments, SH-SY5Y cells were seeded in a 24-wells plate (2×10⁴ cells/cm²) on a coverslip before differentiation process typically 24 h after seeding. Then, cells were fixed with 4% paraformaldehyde for 10 min at room temperature (rt), permeabilized in PBS-Triton X-100 0.1% (PBST) for 10 min at rt and blocked prior to antibody incubation with 5% bovine serum albumin (BSA) in PBS for 45 min at rt. Cells were incubated with primary antibody (rabbit anti-calnexin, Cell Signalling Technology (C5C9) at 1/200 for 1h30 at rt; rabbit anti-RCAS1, Cell Signalling Technology (D2B6N) at 1/200 for 1h30 at rt; rabbit anti-TH, Proteintech (284171AP) at 1/500 for 1h30 at rt; rabbit anti-synaptophysin, Cell Signalling Technology (D8F6H) at 1/200 for 1h30 at rt); and/or direct staining with the anti-β-III-tubulin primary antibody (Thermo Fisher 2G10-TB3 at 1/1000 for 30 min at rt) or with ReadyProbes™ Reagent F-Actin phalloidin red/orange (Invitrogen R37112 at 1/100 for 5 min at rt). The Alexa Fluor 555–conjugated goat anti-rabbit IgG secondary antibody (Life Technologies) was applied at a dilution of 1/10,000. Nuclei were counterstained with DAPI, and samples were mounted using Vectashield (Sigma Aldrich F6057). Fluorescence imaging was performed using a Nikon Eclipse Ti microscope.

### 2.3 RNA extraction

RNA extraction from SH-SY5Y cells (1 well seeded at 2×10⁴ cells/cm²) was performed using the RNeasy Mini Kit (Qiagen 74014) according to the manufacturer’s instructions. RNA was solubilized in 20 µL of RNase-free water.

### 2.4 RT-qPCR

First-strand cDNA was generated by reverse transcription (RT) using the Maxima First Strand cDNA Synthesis Kit for RT-qPCR (Thermo Fisher, K1672) according to the manufacturer’s instructions. The cycling conditions consisted of an initial denaturation at 95 °C for 1 min. Subsequently, 40 amplification cycles were carried out, with denaturation at 95 °C for 15 s and annealing/extension at 60 °C for 1 min, using the C1000 Touch thermocycler (Biorad, CFX96 Real-Time system). Data are expressed as the mean of ratio of 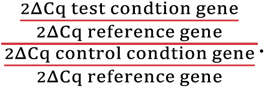 Primers were bought from ThermoFischer: RPLP0 Hs00420895_gH, SYP Hs00300531_m1, TUBB3 Hs00801390_s1, MAP2 Hs00258900_m1, DLG4 Hs01555373_m1, MAPT Hs00902194_m1, RBFOX3 Hs01370654_m1, BCL2 Hs04986394_s1, TFAM Hs00273372_s1, MFN2 Hs00208382_m1, DCX Hs00167057_m1.

### 2.5 MTS and LDH test

Cells were seeded into 96-well plates (2×10⁴ cells/cm²) before differentiation process typically 24 h after seeding. Following treatment with the indicated compounds (H_2_O_2_ from Sigma Aldrich H1009 and Aβ1-42 from Bachem 4014447) for the specified duration, 20 µL of MTS solution (Promega CellTiter 96® AQueous One Solution Cell Proliferation Assay, G3580) was added to each well, and plates were incubated for 1h30. For LDH cytotoxicity test, 50 µL of cell culture supernatants were collected and transferred to a fresh 96-well plate. Then, 50 µL of CytoTox 96® reagent (Promega CytoTox 96® Non-Radioactive Cytotoxicity Assay, G1780) was added according to the manufacturer’s instructions, and the plate was incubated for 30 min at rt protected from light before adding 50 µL of the Stop solution. Plates were stored protected from the light at rt for 30 min before reading. In each case, absorbance was measured at 490 nm using a CLARIOstar Plus plate reader (BMG LABTECH). Cell viability was expressed as a percentage relative to control cells.

### 2.6 Aβ1-42 aggregates preparation

The Aβ1-42 peptide was obtained commercially (Bachem 4014447) and handled according to previously published procedures ^13^. Initially, the peptide was dissolved in 0.16% ammonia in H_2_O (prepared from a 25% ammonia stock solution) at a concentration of 2 mg/mL for aliquoting and was subsequently lyophilized. To disaggregate preformed aggregates, the Aβ1-42 sample was resolubilized in 60 mM NaOH to a final concentration of 1 mg/mL, followed by sonication for 5 min and centrifugation at 5000 rpm for 5 min. The sample was then immediately lyophilized and aliquots were stored at -20 °C until use. Briefly, 20 µg of the monomeric lyophilized peptide was first solubilized in 60 mM HCl prepared in phosphate buffer at 1 µg/µL, then diluted at 50 µM into phosphate buffer (10 mM at pH 7.4) to initiate controlled aggregation. Fibril formation was induced by incubation under conditions favouring self-assembly (37 °C, shaking at 250 rpm, 24 h), while parallel preparations were applied directly to cell cultures in OptiMEM medium (Thermo Fischer 31985062) with 1% FBS and 1% P/S, to assess biological effects at a final 10 µM concentration of Aβ1-42.

## 3. RESULTS

### 3.1 Differentiation protocol and kinetic of differentiation

To induce and characterize the differentiation of SH-SY5Y cells, we established and compared several differentiation protocols. This step was critical to obtain a more mature neuron-like phenotype suitable for downstream functional and molecular analyses. Cells were treated with retinoic acid (RA) and/or brain-derived neurotrophic factor (BDNF) using different treatment durations and combinations (Table 1<u>Table 1</u>). RA is known to promote neuronal commitment and initiate differentiation, whereas BDNF supports neuronal maturation, neurite outgrowth and synaptic functionality ^24^ ^27^. By varying both the combination of these factors and the duration of exposure, we aimed to determine the conditions that most effectively drive the transition toward a stable and functionally mature neuronal phenotype. This comparative approach allowed us to identify differentiation conditions to obtain this neuron-like phenotype and that maximize thereafter reproducibility for subsequent experiments. Structural and transcriptional changes associated with neuronal maturation were assessed throughout the differentiation process.

**Table 1:** Comparison of the protocols tested for SH-SY5Y differentiation. SH-SY5Y cells were exposed to different differentiation conditions using retinoic acid (RA, 10 µM) and/or brain-derived neurotrophic factor (BDNF, 50 ng/mL). The table summarizes the presence or absence of each treatment, the duration of exposure, the starting day of treatment, and the total length of the differentiation protocol for each condition (a–f).

| Conditions | RA<br>(10 μM) | Duration of<br>RA (days) | Starting<br>day | BDNF<br>(50 ng/mL) | Duration of<br>BDNF (days) | Starting<br>day | Total days of<br>treatment |
| --- | --- | --- | --- | --- | --- | --- | --- |
| a | yes | 7 | 1 | no |  |  | 7 |
| b | no |  |  | yes | 7 | 1 | 7 |
| c | yes | 7 | 1 | yes | 7 | 1 | 7 |
| d | yes | 15 | 1 | no |  |  | 15 |
| e | no |  |  | yes | 15 | 1 | 15 |
| f | yes | 15 | 1 | yes | 10 | 5 | 15 |

Immunofluorescence (IF) analyses (Figure 1a) showed that non-differentiated SH-SY5Y cells after 15 days of culture displayed a typical neuroblast-like morphology, characterized by rounded cell bodies and very limited neurite extension. Cells were visualized using F-actin staining, which highlights the cytoskeletal organization and allows clear observation of cell shape and neurite-like outgrowth. Neurite-like processes began to appear as early as 7 days following differentiation with RA alone but this produced relatively modest morphological changes, with only limited neurite-like outgrowth (condition **a**). At this time point, treatment with BDNF alone induced no visible modification of the shape, with cells even less abundant (condition **b**). The effect was slightly more pronounced when RA was combined with BDNF for 7 days (condition **c**), suggesting an enhanced response to the dual treatment; however, these early changes remained partial.

**Figure 1:**
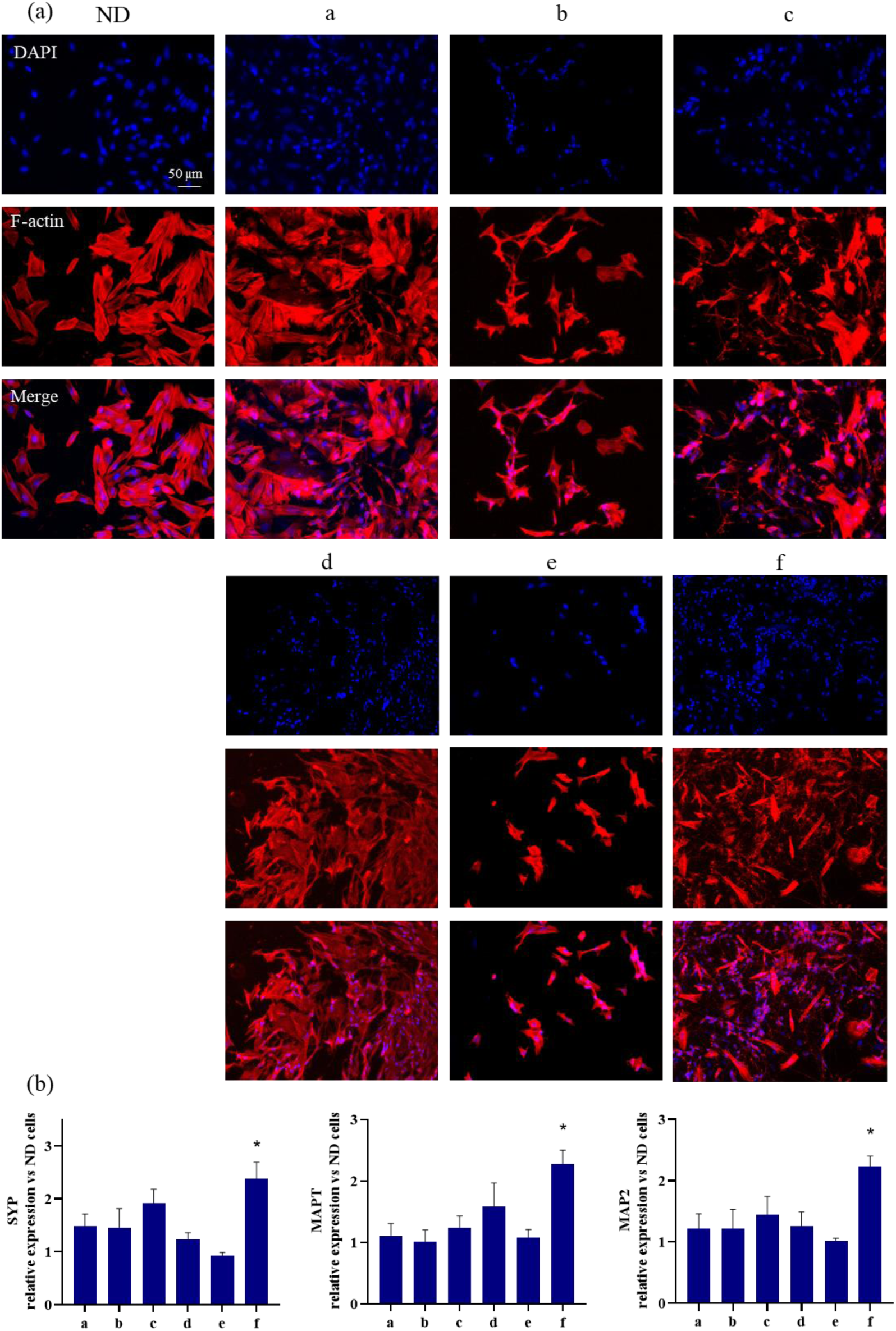
Morphological assessment of SH-SY5Y differentiation under different RA and BDNF treatment protocols. (a) Representative immunofluorescence images of SH-SY5Y cells subjected to the differentiation protocols described before (treatments a-f). Cell nuclei are stained with DAPI (blue) and the actin cytoskeleton is stained with phalloidin red/orange fluorescent conjugate (red). Merge panels show the combined signals. Imaging was performed with a ×10 objective and analysed in the same conditions with ImageJ. (b) Quantification of relative mRNA expression levels measured by RT-qPCR for neuronal markers SYP, MAPT and MAP2 across the different differentiation conditions, compared to the control non-differentiated cells. Data are presented as mean ± SEM. n = 3 independent experiments, each performed in experimental triplicates. Statistical analysis was analyzed using one-sample t-tests against the theoretical value of 1. * *p* < 0.05. ND for non-differentiated cells.

At later time points, clearer morphological alterations were observed. Treatment with RA alone for 15 days (condition **d**) led to the appearance of increased neurite-like extensions, indicating partial differentiation of the cells. In contrast, cells were scarcer and showed minimal neurite-like extension when treated with BDNF alone for 15 days (condition **e**) suggesting that this condition was not favourable for cell health or differentiation. The most pronounced neuron-like morphology was observed when RA and then BDNF were combined, particularly after 15 days of treatment (condition **f**), with increased neurite-like outgrowth and reduced cell body size compared with the 7 days condition, consistent with more advanced differentiation. It should be noted that the nucleus of the differentiated cells is relatively smaller compared to non-differentiated cells.

These structural observations were supported by qPCR analyses using markers specific of neuron-like cells (Figure 1b). Microtubule-associated protein 2 (MAP2) is a cytoskeletal protein primarily localized in dendrites and is widely used as a marker of neuronal structural maturation and dendritic development ^35^. Microtubule-associated protein Tau (MAPT) is involved in the stabilization of microtubules within axons and plays an important role in neurite extension and neuronal polarity ^36^. Synaptophysin (SYP) is an integral membrane protein of synaptic vesicles and is commonly used as a marker of synaptic formation and neuronal connectivity ^37^. Together, the increased expression of these markers reflects both structural and functional aspects of neuronal differentiation.

Conditions **a**, **b**, **c**, **d** and **e** induced no significant variation in the expression of these markers in comparison to non-differentiated cells (Figure 1b). Condition **d** showed a tendency to increase MAPT expression levels but not for SYP and MAP2. Condition **e**, as mentioned in the IF analysis, shows a tendency to a weaker effect on genes expression. In contrast, an increase in the expression of the neuronal differentiation markers SYP, MAPT, and MAP2 was observed in condition **f** compared with non-differentiated cells. Overall, these results were consistent with the morphological observations and further indicated that the differentiation protocol used in condition **f** promotes the most pronounced neuron-like phenotype.

Based on the morphological and transcriptional analyses, the differentiation protocol **f** was selected as the final validated strategy to obtain neuron-like SH-SY5Y cells. In this protocol **f** (Figure 2a), SH-SY5Y cells were first maintained under proliferation conditions and then exposed to RA (10 μM) to initiate neuronal differentiation. RA treatment was applied for 5 days to promote early neurite formation and commitment toward a neuron-like phenotype. Subsequently, cells were co-treated with RA and BDNF (50 ng/mL) until day 15 to support further differentiation. This sequential treatment resulted in pronounced neurite outgrowth, increased neuronal network complexity, and reduced cell body size, features consistent with neuron-like morphology. Fluorescence staining for β-III-tubulin (a neuronal cytoskeleton protein associated with neurite formation) confirmed the neuron-like identity of differentiated cells revealing extensive neuritic processes and interconnected networks (Figure 2<u>Figure 2</u>b). Together, these observations validated the 15-day protocol combining RA priming followed by RA and BDNF co-treatment as the most effective condition to induce robust neuronal differentiation of SH-SY5Y cells.

**Figure 2:**
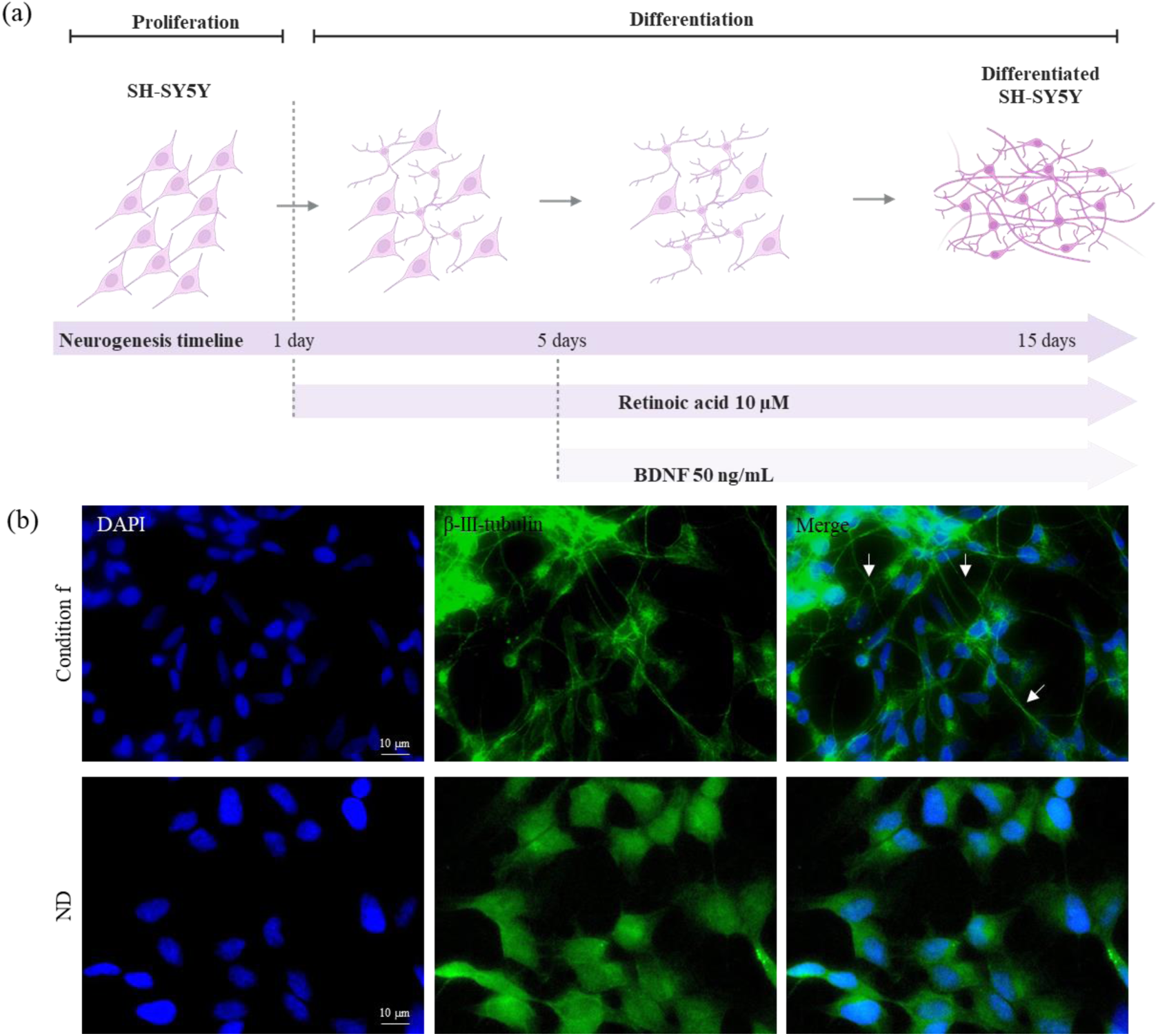
Final protocol of differentiation (protocol f). (a) Scheme representing choosen differentiation protocol. Created with BioRender.com (b) Fluorescence images of SH-SY5Y cells subjected to the differentiation protocol f. Cell nuclei are stained with DAPI (blue) and the β-III-tubulin is detected using specific antibody (green). Merge panels show the combined signals. White arrowheads indicate representative neurite-like extensions. ND for non-differentiated cells. Imaging was performed with a ×60 objective with water imersion.

### 3.2 Triple-level validation of SH-SY5Y differentiation for neuron-like phenotype acquisition

To ensure the reliability of the differentiation strategy (condition **f**), it was important to validate the neuronal phenotype using complementary approaches that assess different aspects of cellular identity and function. Structural changes alone are not sufficient to confirm neuronal differentiation, as neurite-like structures can appear under spontaneous differentiation or stress-related conditions. Therefore, the differentiation protocol was validated using three independent levels of analysis: immunofluorescence to highlight neuronal shape and organelles dynamic, RT-qPCR to quantify the transcriptional upregulation of established neuronal differentiation markers, and functional assays to assess cellular responses relevant to neuronal physiology. This multiparametric validation provided a robust framework to confirm that SH-SY5Y cells acquired a stable neuron-like phenotype under the optimized differentiation conditions.

#### Cell structure validation: cellular shape changed in differentiated SH-SY5Y cells

Immunofluorescence assay was performed to compare differentiated (protocol **f**) and non-differentiated cells by visualizing specific cellular organelles and morphological markers (Figure 3a). As evoked Figure 2b, staining with β-III-tubulin confirmed the extension of neurites in differentiated cells, with a shape drastically different from non-differentiated. Cells were also stained for RCAS1 (Receptor-binding cancer antigene), a protein associated with the Golgi apparatus and involved in protein processing and secretion; for calnexin, an endoplasmic reticulum (ER) membrane chaperone involved in protein folding and quality control; and for synaptophysin. Based on qualitative assessment of fluorescence intensity, a qualitatively moderately stronger RCAS1 labelling was observed in differentiated cells, which may suggest an increased secretory and protein trafficking activity associated with neuronal maturation. Similarly, calnexin labelling seemed moderately increased in differentiated cells, which could be indicative of enhanced ER activity and protein processing during differentiation. In contrast, synaptophysin labelling showed slight difference between differentiated and non-differentiated cells, this protein is already expressed in non-differentiated cells and the differentiation induces slight increase in this expression as we saw in the previous RT-qPCR results (Figure 1b).

**Figure 3:**
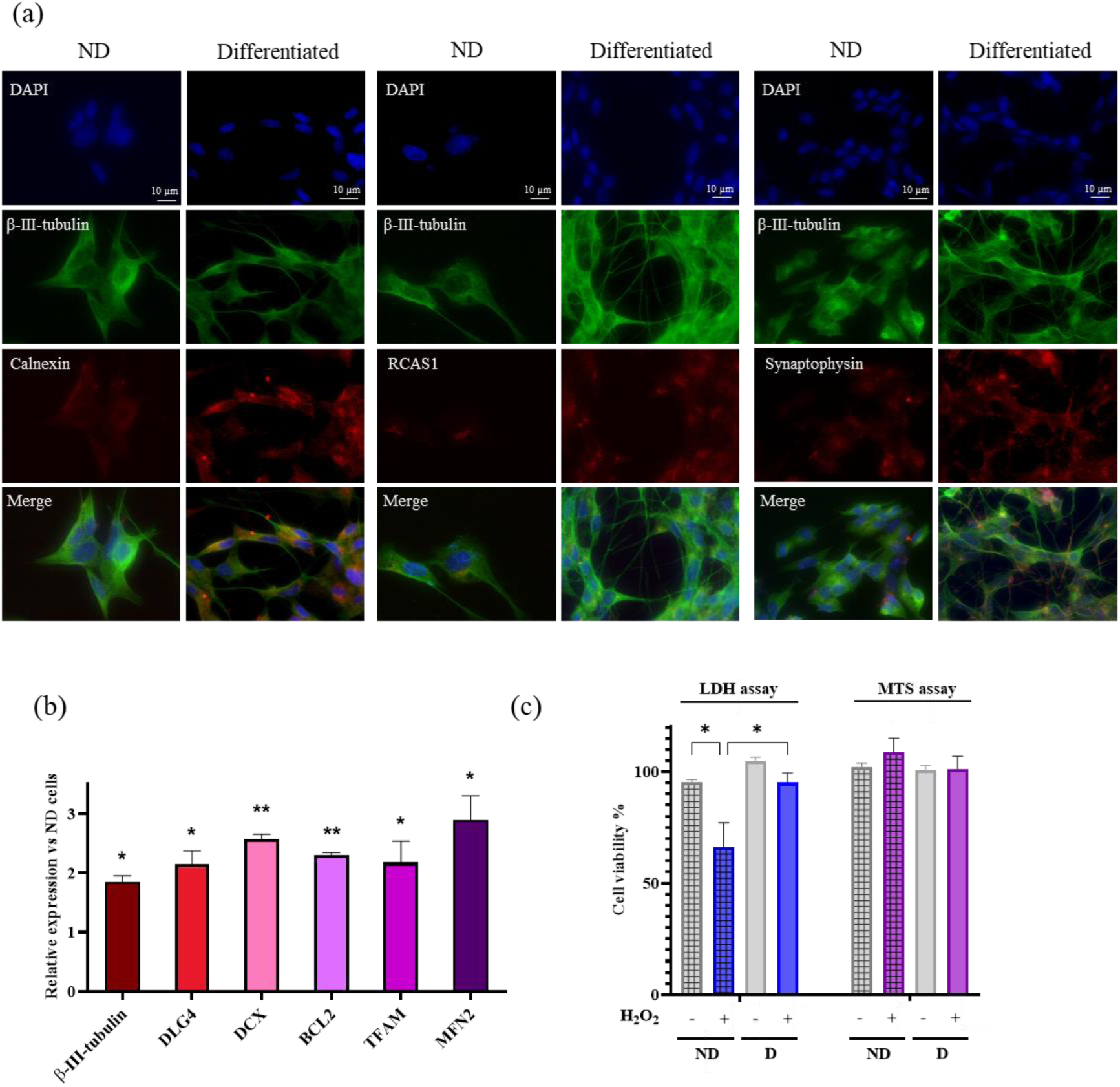
Validation of SH-SY5Y differentiation under protocol. **f.** (a) Representative immunofluorescence images of SH-SY5Y cells subjected to the differentiation protocol f. Cell nuclei are stained with DAPI (blue) and the β-III-tubulin is detected using specific antibody (green), Golgi apparatus (RCAS1), ER (calnexin) and synaptophysin are detected using specific antibody and labelled with secondary antibody (red). Merge panels show the combined signals. Imaging was performed with a ×60 objective with water imersion. (b) Quantification of relative mRNA expression levels measured by RT-qPCR for neuronal markers β-III-tubulin, DLG4, DCX, BCL2, TFAM and MFN2 compared to the control non-differentiated cells (value = 1). (c) Functional test for cell viability assay in presence of H_2_O_2_. Data are presented as mean ± SEM. n = 3 independent experiments, each performed in experimental quadruplicates. Statistical analysis was performed using for (b) one-sample t-tests against the theoretical value of 1 and for (c) two-way ANOVA followed by multiple comparison tests. * *p* < 0.05, ** *p* < 0.01, *** *p* < 0.001, **** *p* < 0.0001. ND for non-differentiated cells. D for differentiated cells.

#### Transcriptional validation: upregulation of selected markers in differentiated SH-SY5Y cells

In addition to the immunofluorescence analyses described above, gene expression profiling was performed as another approach to validate the differentiation protocol (Figure 3b). During the differentiation process, increased expression of several neuronal and cellular markers was observed, including β-III-tubulin, DCX (doublecortin), a microtubule-associated protein involved in neuronal migration and early neuronal differentiation, and DLG4 (disc large homolog 4), a postsynaptic scaffolding protein important for synaptic organization. In addition, the anti-apoptotic gene BCL2, mitochondrial-related genes mitofusin-2 (MFN2), which regulates mitochondrial fusion; and TFAM, a key factor in mitochondrial DNA maintenance and transcription, were also upregulated. Together, these changes supported the acquisition of a neuron-like phenotype and suggest adaptations in mitochondrial function and cellular survival pathways during differentiation.

#### Functional validation: differentiated SH-SY5Y cells were resistant to H_2_O_2_ mediated toxicity

The H_2_O_2_ toxicity test is typically performed as a positive control and model validation step. Hydrogen peroxide is a well-established inducer of oxidative stress, a mechanism strongly implicated in neuronal damage and neurodegenerative processes. By exposing cells to H_2_O_2,_ it is possible to compare the stress sensitivity of differentiated *vs* non-differentiated cells by two distinct viability assays (Figure 3c). The MTS (3-(4,5-dimethylthiazol-2-yl)-5-(3-carboxymethoxyphenyl)-2-(4-sulfophenyl)-2H-tetrazolium) assay was used to assess metabolic activity thus cell viability, and lactate dehydrogenase (LDH) assays measured the LDH release as an indicator of cytotoxicity and cell lysis. The observed increase in LDH release in non-differentiated SH-SY5Y cells indicated that oxidative stress can disrupt plasma membrane in these cells, while the absence of a reduction in MTS signal suggested that mitochondrial metabolic activity was not immediately affected. Conversely, the lack of significant toxicity in differentiated SH-SY5Y cells supported the idea that neuronal differentiation conferred greater resistance to oxidative stress, possibly through improved antioxidant defences or more developed cellular stress-response pathways. This validation step therefore strengthened the relevance of the model before investigating the effects of Aβ1-42 toxicity.

### 3.3 Assessment of Aβ1-42 toxicity in differentiated cells under controlled aggregation kinetic

Assessing Aβ1-42 toxicity in differentiated cells provides a more relevant neuron-like context for investigating its cellular effects. In this study, we aimed at evaluating the effects of both non-aggregated Aβ1-42 (the soluble form) and pre-aggregated Aβ1-42 (the fibrillar form) species. In our previous work, we reported the synthesis of highly pure Aβ1–42 together with a standardized preparation protocol enabling reproducible and controlled aggregation kinetics. This work addressed one of the major, yet often overlooked, sources of variability in amyloid research: the quality and physicochemical state of the peptide itself. ^13^, The aggregation kinetics of Aβ1-42 species were characterized using complementary techniques, including fluorescence spectroscopy, circular dichroism, transmission electron microscopy, dot blot, single-molecule fluorescence imaging and NMR spectroscopy. Together, these methods provided convergent evidence of the progressive structural transitions of Aβ1-42 throughout the aggregation pathway ^13^. Cellular viability and membrane integrity were assessed using MTS and LDH assays. Integrating these measurements in both cell models allowed a comprehensive evaluation of Aβ1-42 toxicity and facilitates the assessment of their respective relevance for studying its mechanism of action.

#### Divergent LDH and MTS responses revealed state-specific Aβ1-42 toxicity in differentiated SH-SY5Y cells

Interestingly, differential responses to Aβ1-42 species were observed between non-differentiated and differentiated SH-SY5Y cells and according to distinct toxicity tests. In this case, Aβ1-42 species were incubated at the end of differentiation. In the LDH assay, non-differentiated cells were predominantly sensitive to pre-aggregated Aβ1-42, both at 48 h and 72 h of incubation, but less sensitive to non-aggregated Aβ1-42 (Figure 4a). Conversely, differentiated cells showed increased cell death following exposure to non-aggregated Aβ1-42, while aggregated Aβ1-42 did not induce significant membrane damage (Figure 4a’), suggesting that soluble Aβ1-42 species had preferential toxicity toward interference with membrane. In the MTS assay, non-differentiated cells displayed minimal reductions in MTS signal, with only a slight decrease observed following soluble Aβ1-42 treatment after 72 h of incubation (Figure 4b). In contrast, differentiated SH-SY5Y cells exhibited reduced metabolic activity following exposure to Aβ1-42 fibrils, indicating mitochondrial dysfunction in response to these species (Figure 4b’). When toxicity was observed, it occurred at least after 48 h of incubation into the cells in both cytotoxicity assays.

**Figure 4:**
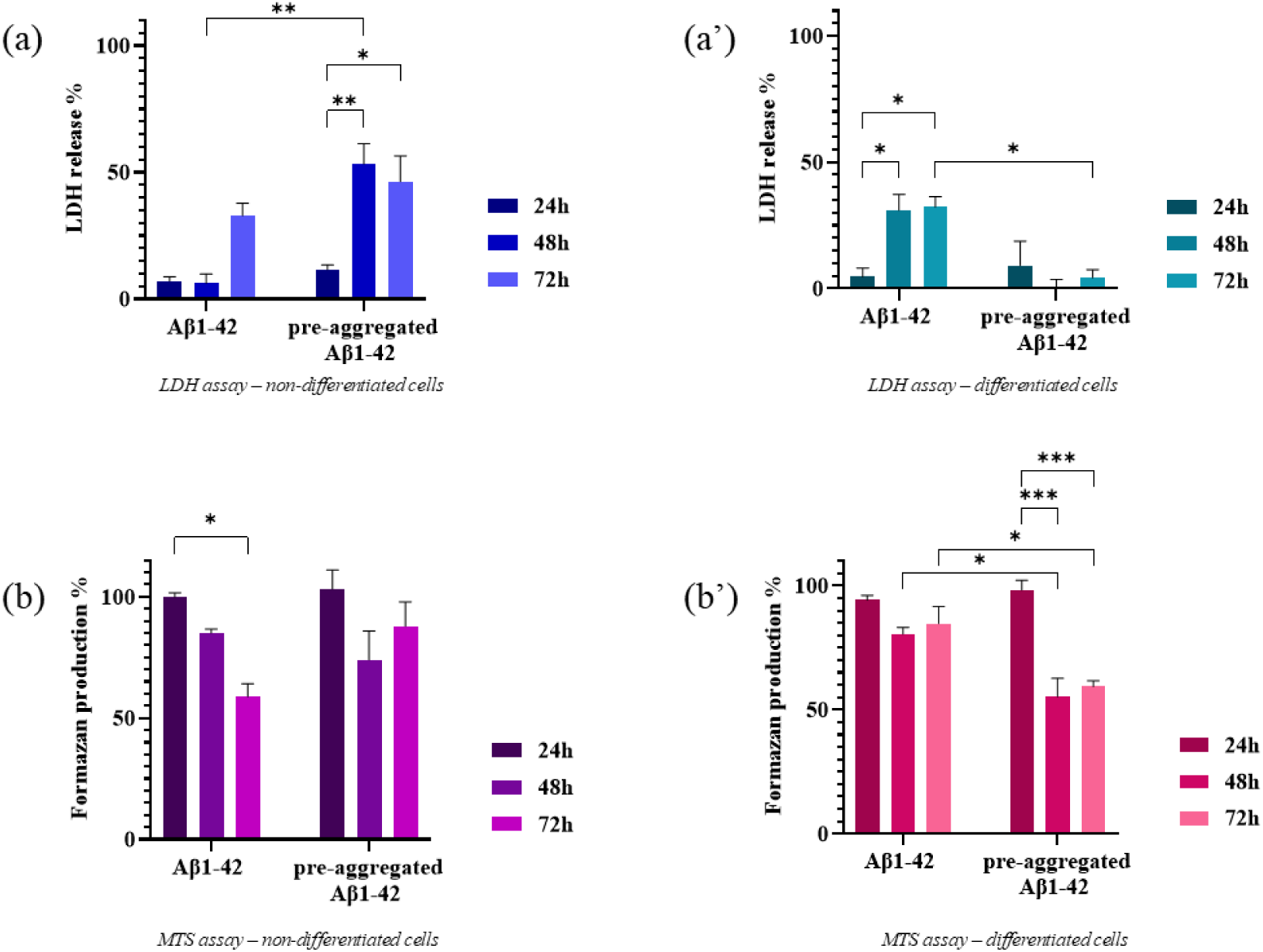
Kinetic of cytotoxic effects of Aβ1-42 pre-aggregated or not on differentiated and non-differentiated SH-SY5Y cells measured by LDH and MTS assays. (a, a’) Cytotoxicity evaluated with LDH test. (b, b’) Cytotoxicity evaluated with MTS test. Aβ1-42 was pre-aggregated or non-aggregated and incubated into the cells for 24 h, 48 h or 72 h. Data are presented as mean ± SEM. n = 3 independent experiments, each performed in experimental quadruplicates. Statistical analysis was performed using two-way ANOVA followed by multiple comparison tests.* *p* < 0.05, ** *p* < 0.01, *** *p* < 0.001.

#### Selective gene expression changes in differentiated SH-SY5Y cells exposed to Aβ1-42

Following the functional toxicity assays, RT-qPCR analysis was performed to assess the molecular responses of differentiated SH-SY5Y cells to Aβ1-42 exposure. Differentiated SH-SY5Y cells were treated with pre-aggregated or non-aggregated Aβ1-42 at the end of differentiation and analysis revealed selective alterations in gene expression with no clear differences between the type of aggregation state (Figure 5<u>Figure 5</u>). Regarding cytoskeletal and synaptic markers, we observed a significant decrease of MAP2 expression for both forms, and β-III-tubulin, MAPT and SYP expression showed a tendency to decrease, suggesting that Aβ1-42 can induce early perturbations in neuronal structural and synaptic components. In contrast, genes related to mitochondrial function and cell survival (TFAM, MFN2, and BCL2) as well as the postsynaptic scaffolding gene DLG4, remained stable, indicating that Aβ1-42 exposure at this stage did not significantly compromise mitochondrial integrity, apoptosis regulation, or postsynaptic organization. Interestingly, the expression of RBFOX3 (NeuN) and DCX highlighted a significant increase after incubation with the both forms and the soluble one respectively, which may reflect compensatory mechanisms or a relative enrichment of cells expressing early neuronal markers in response to Aβ1-42 stress. The observed changes in gene expression primarily reflects early structural or differentiation-related responses rather than overt cytotoxicity. Notably, these findings also suggested that soluble Aβ1-42 induced more pronounced early damage to the cells compared with what was observed in the previous LDH assays.

**Figure 5:**
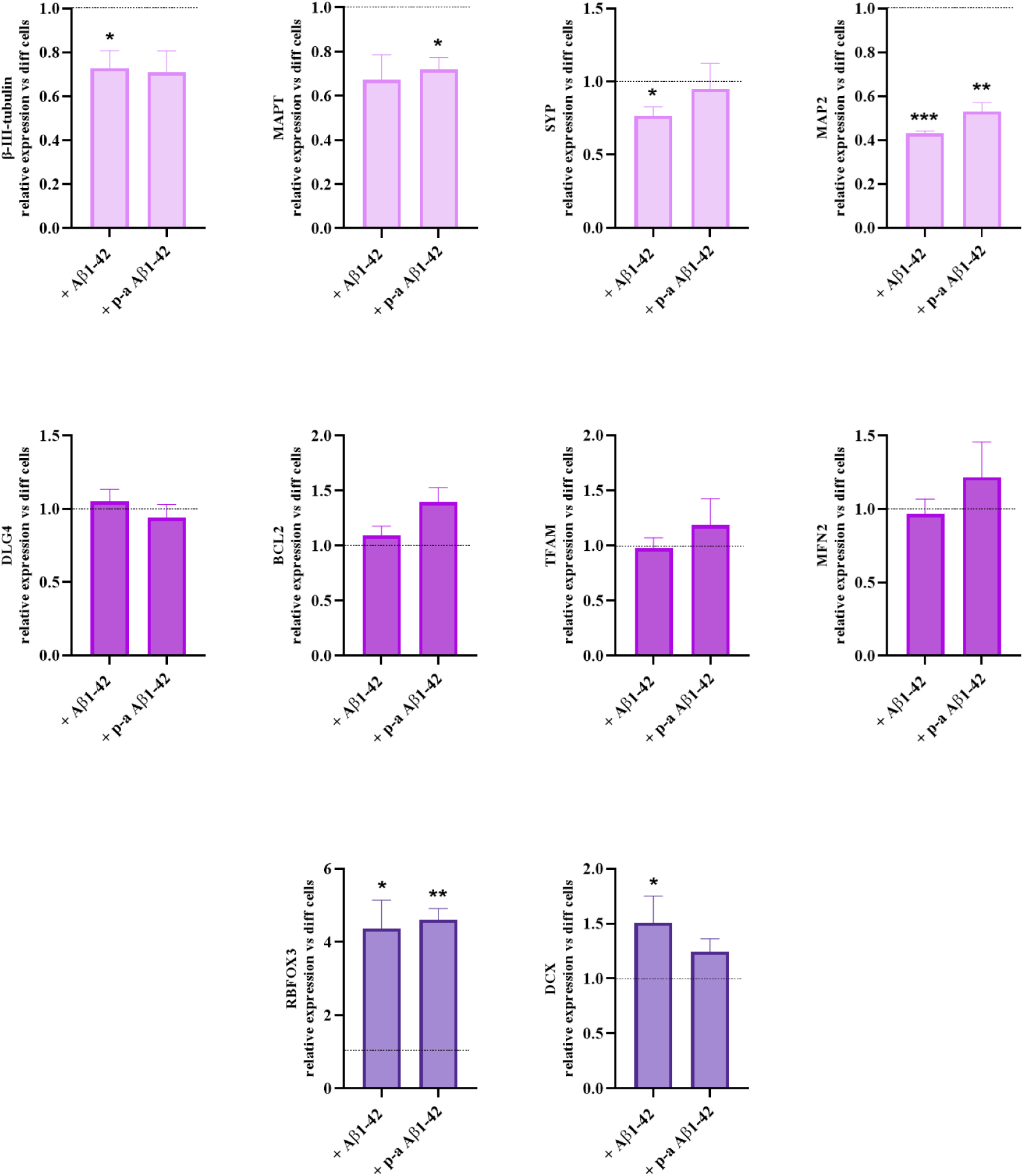
Effect of Aβ1-42 on specific markers. Analysis compared to the control differentiated cells without any treatment (value = 1). p-a for pre-aggregated. Data are presented as mean ± SEM. n = 3 independent experiments, each performed in experimental triplicates. Statistical analysis was performed using one-sample t-tests against the theoretical value of 1. * *p* < 0.05, ** *p* < 0.01, *** *p* < 0.001.

## 4. DISCUSSION

For more than two decades, amyloid-β aggregation inhibitors have been evaluated using experimental workflows that differ considerably among laboratories. Variations in peptide preparation, aggregation protocols, cellular models and toxicity assays have generated a literature that is often difficult to compare and, in many cases, difficult to reproduce.

Although tremendous efforts have been devoted to discovering compounds capable of inhibiting Aβ aggregation, much less attention has been paid to the robustness and standardization of the experimental workflows used to evaluate these molecules.

As a consequence, compounds reported as promising in one laboratory frequently fail to reproduce similar biological effects in another, raising questions not only about the compounds themselves but also about the experimental conditions under which they were evaluated.

In this context, we reported a triply validated neuron-like cell model for the mechanistic and toxicity study of Aβ1-42. Differentiation was confirmed using complementary morphological, transcriptional and functional assays. In parallel, a controlled workflow for the aggregation of Aβ1-42 was developed, providing robust and reproducible framework for studying Aβ1-42 toxicity in a more relevant model ^13^.

At the morphological level, after 7 days of RA treatment, early neurite-like extensions appear, indicating the start of neuronal commitment but with limited growth, and after 7 days of BDNF treatment, almost no transition was observed. A slight improvement with combined RA and BDNF suggests a synergistic effect, although short-term exposure remains insufficient for full differentiation. After 15 days, RA alone induced partial differentiation, whereas BDNF alone failed to promote differentiation. The most robust neuronal phenotype was achieved with the sequential RA and BDNF protocol. This sequential effect reflects distinct but complementary signalling pathways, and reinforce previous reports that RA preconditioning is required for efficient BDNF responsiveness, while further supporting the importance of treatment duration and sequence for achieving robust neuronal phenotypes ^38^ ^39^. qPCR data support these structural observations with increase in MAP2, MAPT and SYP indicating concurrent structural and functional maturation ^40^ ^41^ ^42^. The absence of significant changes in intermediate conditions supports that partial treatments are insufficient to activate the full transcriptional program required for neuronal maturation ^25^ ^43^. Furthermore, as supported by others, non-differentiated SH-SY5Y cells already have a relatively high basal expression of some neuronal markers, so the dynamic range for further induction is limited. Because of this, differentiation tends to produce modest but biologically meaningful changes, rather than large fold increases seen in less specialized systems ^21^ ^24^. However, these moderate changes can still reflect coordinated maturation, especially when multiple markers increase together. The validated 15-days differentiation strategy therefore appears to recapitulate key aspects of neuronal development *in cellulo*. The confirmation of extensive β-III-tubulin-positive neuritic networks further supports the acquisition of a neuron-like identity. Despite these features, SH-SY5Y-derived neurons remain a simplified model that does not fully recapitulate the complexity of primary or iPSC-derived neurons, although they provide a robust and controlled system ^20^ ^44^ ^28^.

Differentiation into a neuron-like phenotype was validated through complementary morphological, transcriptional, and functional analyses, indicating a coordinated maturation process. Morphologically, moderate but consistent changes in organelle-associated markers were observed. However, these observations are based on qualitative assessment of immunofluorescence intensity and were not quantitatively measured. Slight qualitative increased RCAS1 labelling suggests enhanced Golgi activity, consisting with elevated trafficking demands during neuronal maturation ^45^. Similarly, slight qualitative increased of calnexin expression indicates enhanced endoplasmic reticulum activity ^46^. Synaptophysin levels remained relatively stable in agreement with previous reports indicating that SH-SY5Y cells often exhibit incomplete synaptic maturation and its basal expression in undifferentiated cells ^21^. This observation aligns with prior qPCR results showing only modest transcriptional upregulation, suggesting that while synaptic components begin to emerge, full synaptic maturation may not yet be achieved under the applied differentiation conditions. Transcriptional profiling further supports the acquisition of neuron-like characteristics. The upregulation of β-III-tubulin and DCX confirms cytoskeletal remodelling and early neuronal differentiation ^47^ ^48^, while upregulation of DLG4, coding for PSD95 a key postsynaptic scaffolding protein, indicates the initiation of synaptic organization, consistent with studies showing early synaptic gene induction preceding full synaptogenesis ^49^. Differentiation was also associated with enhanced cellular resilience. Elevated expression of BCL2 is consistent with its role in survival during neuronal maturation and increased expression of MFN2 and TFAM reflects mitochondrial adaptations critical for neuronal bioenergetics ^50^ ^51^. These changes are particularly relevant given the high demands of neurons and the central role of mitochondria in supporting neuronal activity and survival ^52^. Functionally, differentiated cells exhibited increased resistance to H_2_O_2_-induced stress. While non-differentiated SH-SY5Y cells showed increased LDH release following oxidative challenge, differentiated cells maintained both membrane integrity and metabolic activity, indicating a more robust stress response. This increased resilience is consistent with enhanced antioxidant defences, stress-response pathways, and upregulation of BCL2, along with mitochondrial adaptations involving MFN2 and TFAM ^24^ ^25^ ^52^. Additionally, mitochondrial adaptations reflected by increased MFN2 and TFAM expression may contribute to maintaining cellular homeostasis under stress conditions. Overall, these findings demonstrate that SH-SY5Y differentiation induces coordinated changes in organelle function, gene expression, and stress response, supporting the acquisition of a more neuron-like phenotype. This highlights the fact that not all cell models are equivalent, and that a toxic response can vary depending on the cells stage of differentiation.

In this context, this validated system therefore provides a robust platform for studying neuronal function and vulnerability, particularly in the context of neurodegenerative processes such as Aβ1-42 toxicity. The present study demonstrates that the toxicity of Aβ1-42 in SH-SY5Y cells is strongly dependent on (i) the aggregation state, (ii) the differentiation state of the cells and (iii) the assay used to assess cell viability. The combined use of LDH and MTS assays provides important insight into the temporal and mechanistic aspects of Aβ1-42 aggregates toxicity. In this study, LDH assays revealed that differentiated cells (but not non-differentiated cells) are more sensitive to non-aggregated Aβ1-42, highlighting the membrane-disruptive properties of these species, when fibrillar Aβ1-42 did not induce significant membrane damage. In contrast, MTS measurements showed higher impact of fibrillar species on these differentiated cells (but not on non-differentiated cells). This divergence underscores the importance of assay selection when interpreting cytotoxic effects, as different readouts capture distinct stages of cellular stress. The apparent discrepancies between LDH and MTS results reflect the distinct cellular processes measured by these assays, suggesting that Aβ1-42 toxicity in SH-SY5Y cells manifests through distinct cellular mechanisms depending on both differentiation status and peptide aggregation state.

At the same time, the modest alterations observed in structural and synaptic markers suggest that Aβ1-42 primarily induces subtle, early-stage toxicity rather than overt neurodegeneration in this model. Interestingly, exposure to Aβ1-42 resulted in a marked increase in RBFOX3 expression, accompanied by a modest upregulation of DCX, despite the presence of functional toxicity. Interestingly, it suggests that Aβ1-42 does not immediately suppress neuronal identity but instead triggers adaptive cellular responses. RBFOX3 upregulation may reflect a stress-induced adaptation in RNA processing, while increased DCX expression suggests activation of cytoskeletal remodelling and plasticity-related pathways. Importantly, the absence of significant changes in mitochondrial (TFAM, MFN2) and anti-apoptotic (BCL2) gene expression indicates that these responses occur independently of major transcriptional reprogramming of survival or metabolic pathways. Together, these findings suggest that Aβ1-42 does not markedly alter the transcription of key mitochondrial and survival genes under the tested conditions, and that its exposure induces an early adaptive phase characterized by reinforcement of neuronal identity and structural plasticity, preceding the activation of irreversible degenerative processes. Aβ1-42 toxicity is not purely degenerative, it also triggers adaptative neuronal responses and this depends on peptide aggregation and cells differentiation states. Basically, in this system, there is a first phase highlighted by adaptive neuronal stress response, followed by functional impairment leading to assumingly a last apoptotic phase that we have not reached yet.

Differentiated SH-SY5Y cells represent an intermediate neuron-like state characterized by partial maturation, intrinsic neuronal features, and increased resistance to apoptosis. This profile makes them particularly suitable for modelling early stages of neurodegenerative processes, where subtle functional and structural alterations precede overt cell death. In this context, exposure to Aβ1-42 induces mild but detectable changes in neuronal integrity, especially in response to soluble species, while largely preserving mitochondrial function and viability under the tested conditions. These findings reinforce the relevance of this model for investigating early pathogenic events associated with Aβ1-42 toxicity. More broadly, they highlight that cellular responses to Aβ1-42 are highly context-dependent, varying with differentiation status and methodological readouts, which may contribute to the variability observed across studies. Altogether, this work emphasizes the importance of combining well-characterized models with complementary assays to accurately capture the complexity of Aβ1-42-induced neurotoxicity.

## RESOURCE AVAILABILITY

Lead contact: Further information and requests for resources and reagents should be directed to the lead contact, Nicolo Tonali.

Materials availability: All unique/stable reagents generated in this study are available from the lead contact without restriction.

Data and code availability • All data reported in this paper will be shared by the lead contact upon request. • This paper does not report original code. • Any additional information required to reanalyze the data reported in this paper is available from the lead contact upon request.

## ACKNOWLEDGMENT

We are grateful to many participants, researchers and staff who contributed significantly to this study (Marion Chaigneau, Peggy Barbe). This work was supported by the Agence Nationale de la Recherche and the Deutsche Forschungsgemeinschaft (DFG) through the joint ANR–DFG program (ANR-22-CE92-0076-01, DFG SE 609/20-1), as well as by PASREL (Université Paris-Saclay), France 2030 programme (ANR-11-IDEX-003) and by BIOPROBE (Université Paris-Saclay), France 2030 programme (ANR-11-IDEX-003).

## AUTHORS CONTRIBUTION

A.V. and N.T. conceived the project. A.V. and C.P. performed the experiments. A.V. and C.P. analyzed the data. A.V. wrote the manuscript. X.I., P.R., S.P., N.S., D.S., N.T. edited and revised the manuscript.

## CONSENT STATEMENT

Consent was not necessary.

## ABBREVIATIONS

BCL2: B-cell lymphoma 2
BDNF: brain-derived neurotrophic factor
DCX: doublecortin
DLG4: disc large homolog 4
ER: endoplasmic reticulum
ESC: embryonic stem cell
FBS: fetal bovine serum
HCl: hydrochloric acid
iPSC: induced pluripotent stem cell
LDH: lactate deshydrogenase
MAP2: microtubule-associated protein 2
MAPT: microtubule-associated protein tau
MFN2: mitofusin-2
NMR: nuclear magnetic resonance
MTS: 3-(4,5-dimethylthiazol-2-yl)-5-(3-carboxymethoxyphenyl)-2-(4-sulfophenyl)-2H-tetrazolium
NaOH: sodium hydroxide
P/S: penicillin/streptomycin
PBS: phosphate buffer saline
RA: retinoic acid
RCAS1: receptor-binding cancer antigene
RT-qPCR: reverse transcription-quantitative polymerase chain reaction
SYP: synaptophysin
TFAM: mitochondrial transcription factor A

